# Direct Arp2/3-vinculin binding is essential for cell spreading, but only on compliant substrates and in 3D

**DOI:** 10.1101/756718

**Authors:** Tadamoto Isogai, Kevin M. Dean, Philippe Roudot, Qiuyan Shao, Justin D. Cillay, Erik. S. Welf, Meghan K. Driscoll, Shaina P. Royer, Nikhil Mittal, Bo-Jui Chang, Sangyoon J. Han, Reto Fiolka, Gaudenz Danuser

**Affiliations:** Lyda Hill Department of Bioinformatics, University of Texas Southwestern Medical Center. Dallas, TX, USA; Department of Cell Biology, University of Texas Southwestern Medical Center. Dallas, TX, USA; Department of Biomedical Engineering, Michigan Technological University. Houghton, MI, USA

## Abstract

Cells modify their shape in response to the extracellular environment through dynamic remodeling of the actin cytoskeleton by actin-binding proteins (ABPs) ^1–4^. The relation between actin dynamics and spreading is well-understood for cells on flat glass coverslips; much less is known about cell morphogenesis in compliant three-dimensional environments, and, in particular, how ABPs contribute to this process ^5^. Here, we knocked-out a diverse set of ABPs, and evaluated the effect of each on cell spreading on planar glass surfaces (2D) and in reconstituted collagen gels (3D). Our morphometric analyses identify the Arp2/3 complex and its associated regulatory genes among the ABPs that contribute most strongly to cell spreading in 3D, but marginally in 2D. Cells lacking Arp3 have reduced spreading specifically in 3D, and display stiffness-dependent cell-matrix adhesion defects. Through manipulation of vinculin activity, we determine that the Arp3 knock-out phenotype largely arises from the lack of direct interaction between vinculin and Arp2/3 complex. This interaction is dispensable for cell spreading in 2D. These data highlight that actin architectural features necessary for adhesion formation and cell spreading in 3D are efficiently compensated on flat and stiff surfaces.

---

Cell shape control is impacted by extracellular matrix (ECM) mechanics, composition, and architecture ^1,2^. Most of our understanding of the mechanism of cell morphogenesis stems from experiments performed on flat and stiff environments such as glass or plastic. On these planar surfaces (2D), cells are able to spread without restrictions and usually maximally stretch themselves, resulting in a flat morphology with abundant filamentous actin (F-actin) stress fibers (Fig. 1a). When the same cells are embedded in a three-dimensional reconstituted matrix (3D), such as collagen gels, they adopt a multipolar branched morphology with diminished stress fiber formation ^6–9^ (Fig. 1b,c). These differences are in part manifestation of an altered organization of the F-actin cytoskeleton, which is governed by differential association and activation of ABPs in response to shifts in the cell environment ^3,10–14^.

**Figure 1.**
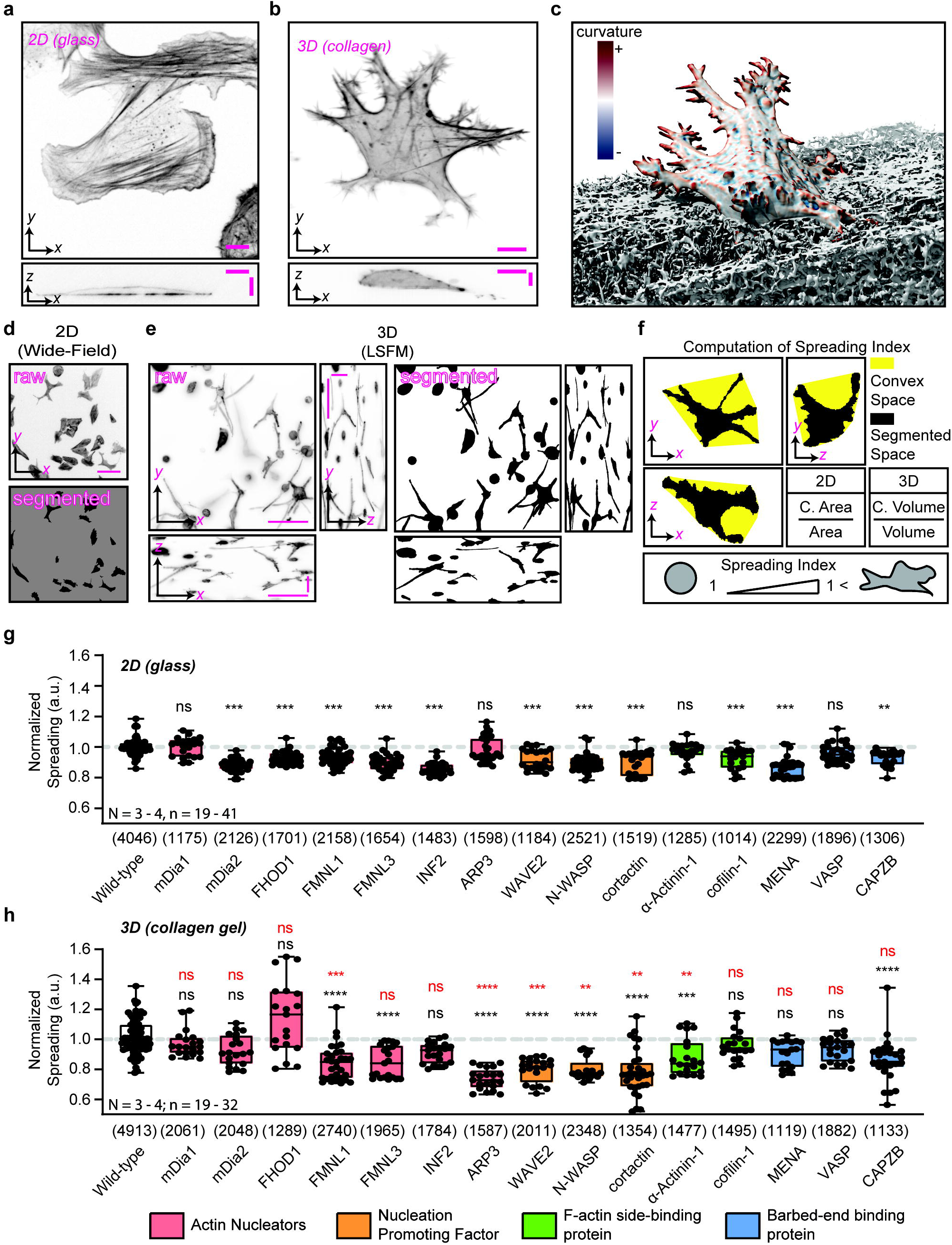
Actin Binding Proteins regulate cell spreading differentially in cells cultured on 2D substrates in 3D matrix. **a**, U2OS cells display a flat morphology on thin (2 µg/ml) collagen-coated glass coverslips **b**, U2OS cells embedded in reconstituted collagen gels (3 mg/ml) display complex mixture of branched protrusions. **a,b**, Cells were stained with fluorescent phalloidin and images were taken using ASLM ^39^, and are representative from three experiments. Scale bars, 10 µm. **c**, Cells rendered in 3D using ChimeraX with the collagen environment partially removed to reveal the cell morphology ^40^. **d,e** Examples of cell segmentations in **d** 2D and **e** 3D (shown as maximal intensity projections (MIP). Scale bars, 100 µm. **f**, Computation of spreading index defined by the ratio between convex space and segmented space (space: area for 2D and volume for 3D). Cells with a spreading index of 1 are convex and lack detectable protrusions. The index monotonically increases as cells become more protrusive and branched. **g,h**, Quantitation of cell spreading in **g** 2D and **h** 3D. All statistical testing was performed in comparison to the wild-type (black asterisks). Statistical comparison of the distribution of spreading indices between 3D and 2D are denoted with red asterisks. All data were collected from at least three independent experiments (N). Each data point represents the median of the spreading indices of all cells within one image field of view (n), and were each normalized to wild-type control. Total number of quantified individual cells are indicated in brackets. Boxplots: Min/Max. Kruskal-Wallis test: ns = not significant, ** p ≤ 0.01, *** p ≤ 0.001, **** p ≤ 0.0001.

To gain insight into the roles of diverse ABPs play in supporting 2D versus 3D cell morphologies, we performed a targeted CRISPR/Cas9-mediated knock-out (KO) screen of formins (mDia1/*DIAPH1*, mDia2/*DIAPH3*, FHOD1/*FHOD1*, FMNL1/*FMNL1*, FMNL3/*FMNL3*, INF2/*INF2*) ^15^, the Arp2/3 complex subunit (Arp3/*ACTR3*) ^14^, Arp2/3 complex-activating Nucleation Promoting Factors (NPFs: Wave2/*WASF2*, N-WASP/*WASL*, cortactin/*CTTN*) ^13^, F-actin side-binding proteins (α-Actinin-1/*ACTN1*, cofilin-1/*CFL1*) ^16^, and barbed-end regulators (CapZβ/*CAPZB*, MENA/*ENAH*, VASP/*VASP*) ^17^. None of the KOs caused noticeable growth defects under standard culture conditions, and all KOs were confirmed at the protein level (Supplementary Fig. 1).

We first measured cell spreading on planar surfaces. Cells were seeded on thin collagen-coated (2 µg/ml) glass coverslips and allowed to adhere and spread for 16 hours. To quantify the differences between cell morphologies of different knock-outs, we segmented individual cells based on fluorescent F-actin staining and computed the ratio between the area of the convex hull embedding the entire cell perimeter and the segmented cell area as an overall index of cell spreading (Fig. 1d,f). Cells with a non-polarized round morphology will have a spreading index close to 1. As cells polarize and extend protrusions the index monotonically increases without bounds (Fig. 1f). These quantitative analyses revealed that most knock-outs except those of mDia1, Arp3, α-Actinin-1 and VASP caused significant, yet minor spreading defects (Fig. 1g).

Strikingly, the subtle defects of ABP KOs in 2D translated into pronounced defects when cells were placed in reconstituted collagen I gels (3D; 3 mg/ml; Fig. 1h). In these experiments, cells were allowed to spread for 16 hours prior to fixation, F-actin staining, and imaging. Consistent with the 2D screen, we segmented individual cells in the acquired 3D image stacks and computed the ratio between convex volume and the segmented cell volume as the spreading index in 3D (Fig. 1e,f).

Using the index, we discovered that KOs of mDia2, INF2, cofilin-1, and MENA, which showed some spreading defects in 2D (Fig. 1g), did not affect cell spreading in 3D (Fig. 1h). Moreover, as in 2D, mDia1 and VASP KOs showed no effects (Fig. 1h). Amongst the tested formins, both FMNL1 and FMNL3 KOs displayed spreading defects in 2D and 3D. The defect of FMNL1 KO was larger in 3D than in 2D (red asterisks in Fig. 1h), suggesting that FMNL1 may have distinct contributions to cell spreading specifically in 3D. Overall, on 2D, most ABP KOs exhibited some degree of spreading defects, although the defects in 3D tended to be more severe (Fig. 1h). This suggests that anomalies in the ABP functions are more tolerated in 2D, possibly due to compensatory actions between ABPs with partially overlapping functions ^18^.

The strongest amplification of spreading defects when switching from 2D to 3D was observed for KOs targeting the Arp2/3 complex-associated dendritic network machinery, including Arp3, Wave2, N-WASP and cortactin (Fig. 1h). Most remarkably, while Arp3 KO cells were not affected in 2D (Fig. 1g), these same cells showed the strongest decrease in the spreading index of all tested KO conditions (Fig. 1h). Moreover, in 2D, these cells displayed a heterogeneous response, reflected by a broad distribution in the spreading index. The same cells in 3D were highly consistent and displayed a tight distribution (for n >1500 cells). Again, we interpreted this outcome as a result of compensation by other ABPs overcoming Arp2/3 depletion in 2D, whereas in 3D, Arp2/3 assumes a specific function that is essential for spreading.

Arp2/3 is the only nucleator of branched actin. Hence, we hypothesized that dendritic actin networks serve a special function during 3D cell spreading. To examine this, we first validated the 3D cell spreading defects under Arp3 knockout by generating additional KO cells line with an orthogonal guide RNAs (gRNAs; Fig. 2a,b). The gRNA targeting Arp3 employed in the screen (Fig. 1g,h) and the gRNA to generate the second Arp3 KO cells lines (Fig. 2a) both targeted the first exon from the *ACTR3* gene ^19^, and resulted in Arp3 variant 1-specific KOs (Arp3 KO #1 and #2; Supplementary Fig. 2a). Arp3 variant 2 ^19^ has an alternative first exon and is unaffected by these gRNAs (Supplementary. Fig. 2). Therefore, we designed a third gRNA that targets both transcript variants, which completely ablates Arp3 expression (Arp3 full KO), and these cells also had similar 3D spreading defects as the variant specific KOs (Fig. 2a,b). The heterogeneous distribution of cell spreading indices of the Arp3 KO #2 (Fig.2b) is likely due to incomplete KO (Supplementary Fig. 2a). The Arp3 KO #1 and Arp 3 full KO were accompanied by degradation of the ARPC2 subunit (Supplementary Fig. 2a), in agreement with previous studies ^20^. This suggests that spreading defects caused by Arp3 loss are likely due to decreased Arp2/3 complex activity, as ARPC2 facilitates the binding of the complex to F-actin ^14,21^. Indeed, acute inhibition of the complex activity using the inhibitor CK-666 ^22^ phenocopied both the lack of phenotype in 2D and the cell spreading defects in 3D of Arp3 KO cells (Fig. 2c,d; Supplementary Fig. 2b,c). Thus, we attribute the phenotype of Arp3 KO to decreased nucleation of dendritic network.

**Figure 2.**
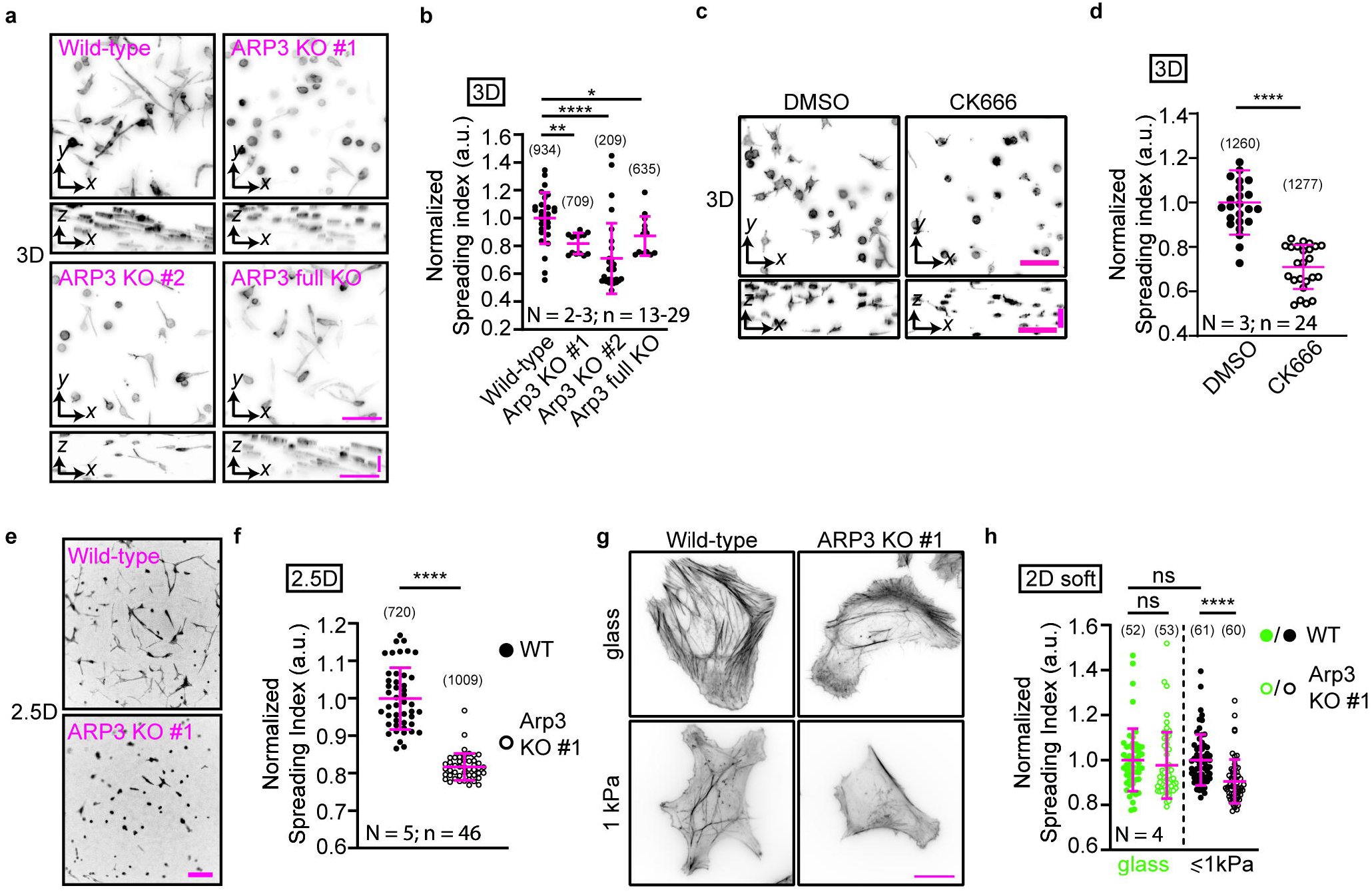
Arp2/3 complex-nucleated dendritic actin network is responsive to ECM dimensionality and stiffness and required for cell spreading in soft ECM environments. **a,b**, U2OS cells lacking Arp3 variant 1 (Arp3 KO #1 and #2) or variants 1 and 2 (Arp3 full KO), have spreading defects in 3D. Cells embedded in collagen gels were stained for F-actin, imaged and quantified as described in Methods. Spreading Index is normalized to wild-type control. **c,d** Polymerization of the dendritic actin network is required for cell spreading in 3D. Collagen-embedded cells were treated overnight with Arp2/3 inhibitor CK-666. Cells were analyzed as in a,b and normalized to DMSO control. **e,f**, Arp2/3 complex is required for cell spreading on soft 2D collagen gels. Cells were seeded on pre-polymerized collagen gel (3mg/ml) and the spreading indices were computed as in b. Data is normalized to wild-type control. **g,h**, Arp2/3 complex is required for cell spreading on soft substrates. Cells were seeded on soft silicone substrates ranging from 0.7 – 1.0 kPa in stiffness and spreading was quantified. **b,d,f,h**, All data were collected from two to five independent experiments (N). Each data point represents the median of the spreading indices of all cells within one image field of view (n), except for h, where each point represents a single cell. Total number of individual cells are indicated in brackets. Data are shown as mean ± s.d. Kruskal-Wallis test: ns = not significant, * p ≤ 0.05, ** p ≤ 0.01, *** p ≤ 0.001, **** p ≤ 0.0001. Scale bars are 100 µm for **a,c,e** and 20 µm for **g**.

The collagen gels used for 3D culture are both geometrically and mechanically different than glass substrates. Geometrically, cells on glass can spread in any direction and form integrin-mediated adhesions without physical impediment, whereas cells in 3D must spread within a confined space and with the restriction that adhesions can form only in proximity to the collagen fibrils. Mechanically, collagen gels are softer than collagen-coated glass (~0.2 – 0.6 kPa vs >MPa ^7,23^). Thus, we asked whether the fibrillar 3D geometry or the stiffness of the extracellular environment turn Arp3 into an essential gene for cell spreading in 3D.

To address this, we seeded wild-type and Arp3 KO cells on top of a pre-polymerized collagen gel (2.5D), altering the confined geometry while minimally changing the stiffness and fibrillar architecture of the ECM. Under these conditions, Arp3 KO cells continued to display a spreading defect relative to the wild-type cells (Fig. 2e,f). Hence, the increased compliance and the fibrillar architecture of the ECM in 2.5D appeared to be sufficient to reproduce the spreading defects of Arp3 KO cells in 3D.

To further test whether the fibrillar collagen architecture is the key to impede cell spreading of Arp3 KO cells, we used flat silicone-based substrates with defined stiffnesses (0.7 – 1 kPa). These substrates resemble a surface similar to that of 2D glass samples lacking collagen fibrils (Fig. 1g), but with the stiffness of collagen gels. On these substrates, the Arp3 KO cells consistently displayed reduced spreading compared to the wild-type cells (Fig. 2g,h). However, the overall reduction of spreading was less compared to that of cells seeded on 2.5D or in 3D (Fig. 2b,f). Thus, these results suggest that architectural features in addition to the stiffness of the ECM attenuate cell spreading of Arp3 KO cells.

In view of the spreading defects of Arp3 KO cells on soft substrates, we hypothesized that the Arp2/3 function relates to cell adhesion, possibly via nucleation of a dense branched actin network that supports the formation and regulation of nascent and focal adhesions (NAs and FAs, respectively) in response to stiffness ^20,24^. To test this, we imaged the adhesion protein paxillin in wild-type and Arp3 KO cells with Total Internal Reflection Fluorescence-compatible 2D soft substrates ^25^ and used previously described computer software to distinguish NAs from mature FAs ^26^. These analyses showed that, although statistically insignificant, Arp3 KO cells had less NAs per cell area on stiff glass substrates compared to WT cells. This difference became significant on increasingly softer substrates (Fig. 3a-c). In contrast, Arp3 KO cells showed similar levels of mature FAs, with only a marginal decrease at lower stiffnesses (Fig. 3a-c). Thus, Arp3 seems to specifically contribute to NA formation in response to substrate stiffness.

**Figure 3.**
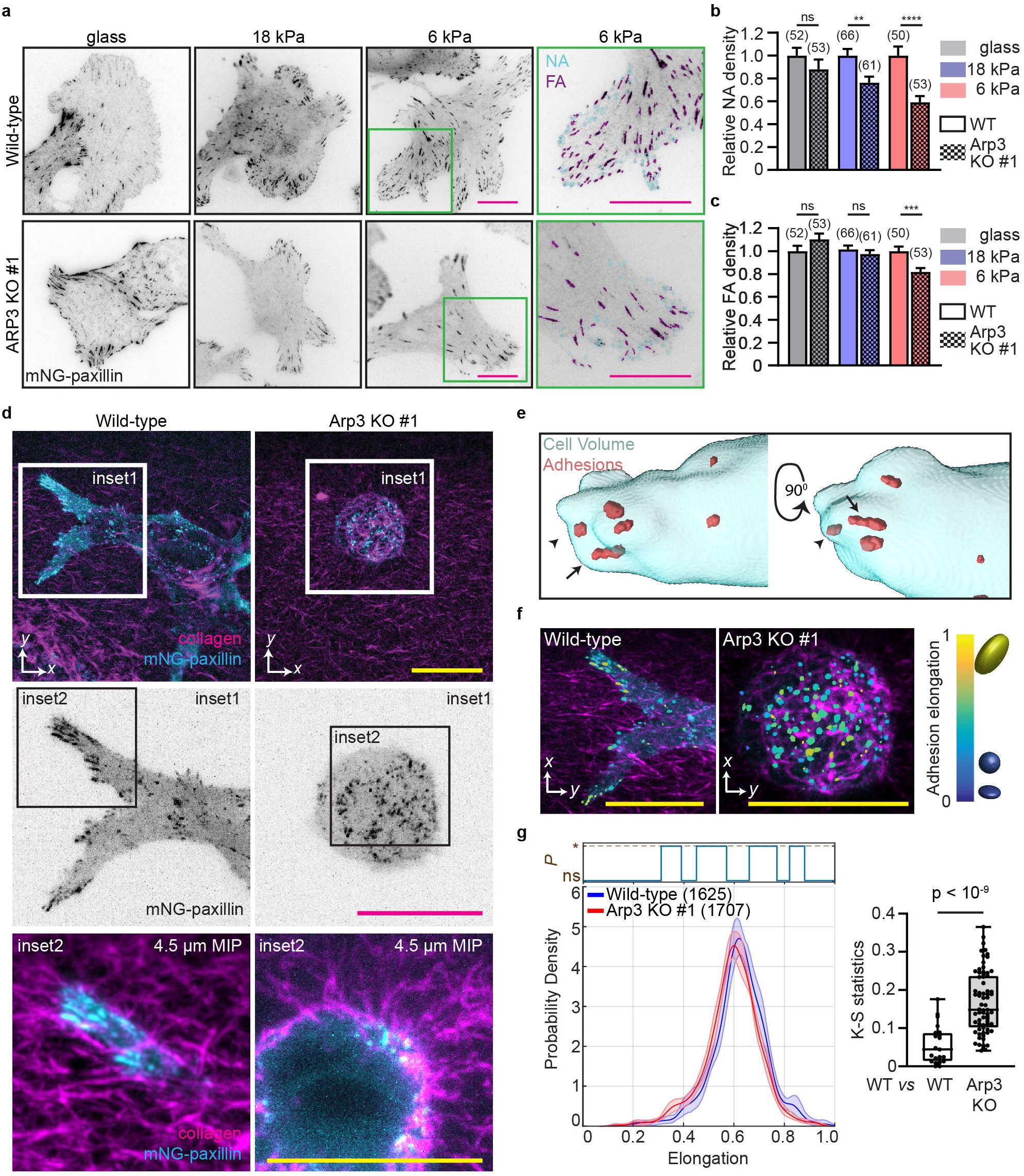
Arp2/3 complex regulates cell-matrix adhesions in a stiffness-dependent manner. **a**, Focal adhesions in cells seeded on collagen-coated glass or silicone substrates were imaged using TIRF. Adhesions, visualized with mNeonGreen-paxillin were classified into nascent adhesions (NA) or mature focal adhesions (FA) using a previously described software ^27^ (Methods). **b,c**, Differential effects of Arp3 KO on NA and FA density, with increasing sensitivity on softer substrates. **b** NA and **c** FA densities (number/cell area) were quantified using a previously described software ^26^ (Methods). Data were collected from four independent experiments (N), and shown as mean ± s.e.m. Number of individual cells are indicated in brackets. t-test: ns = not significant, ** p ≤ 0.01, *** p ≤ 0.001, **** p ≤ 0.0001. **d**, Arp3 KO cells have adhesion defects in 3D. MIPs of adhesions (cyan; mNeonGreen-paxillin) of cells embedded in alexa568-labeled collagen (magenta) are shown. Inset1 illustrates elongated and non-elongated adhesions. Inset2 shows a 4.5 µm MIP of protruding areas of a cell. **e**, 3D rendering examples of elongated (arrow) and non-elongated (arrowhead) focal adhesions in 3D. **f**, Arp3 KO cells have reduced elongated adhesions in 3D. The degree of elongation of adhesions in three-dimensions were quantified with a customized 3D tracking software (Methods). Adhesions with an elongation index of 0 are fully round and increases as they become stretched. Adhesions are pseudocolored according to the elongation index. **g**, *left*, Probability density distribution of adhesion elongation index. Shaded areas represent 95% confidence intervals of the mean adhesion elongation. Statistical significance (*P*) (ns = not significant; * = p < 0.05) of the difference between the elongation index distributions is assessed for 100 non-overlapping 1% quantiles in the range of [0 … 1], is determined by t-test. *right*, Distributions of K-S statistics of the difference in elongation index distributions between either individual wild-type (WT) cells or between WT and Arp3 KO cells (7 cells for WT and 10 cells for Arp3 KO from a representative experiment out of two independent experiments) further confirms the significance of adhesion elongation differences between these two conditions. Total of 1625 and 1707 individual adhesions for WT and Arp3 KO cells, respectively were analyzed, and are indicated in brackets. All scale bars, 20 µm.

We then hypothesized that the cell spreading deficiency observed for Arp3 KO cells in 3D would also arise from a defect in adhesion formation. We evaluated this possibility by imaging paxillin in collagen-embedded cells. Because wild-type cells spread more than Arp3 KO cells and occupy a greater volume, the NA density per area or volume for Arp3 KO cells is inflated (Fig. 3d). Instead, using customized tracking software ^27^ (Methods), we measured the elongation of adhesion-like structures as a proxy for the level of engagement of an adhesion with a collagen fiber (Fig. 3d-g). These analyses showed that wild-type cells have significantly more elongated adhesion-like structures than Arp3 KO cells (Fig. 3f,g), again pointing at a defect in cell-substrate interactions under Arp3 KO conditions. While in 2D, defects in Arp3-dependent adhesion formation pathways are compensated by alternative pathways in an already overall saturated density of adhesions, in 3D, where adhesion formation is rate-limited by the fibrillar topology of extracellular ligands ^6,28^ and is responsible for balancing intracellular contractility in all dimension, such defects translate immediately into diminished spreading.

Next, we considered the underlying molecular mechanisms for Arp3 in adhesion formation. There are two mutually non-exclusive possibilities: First, there is a direct requirement for the presence of the Arp2/3 complex in the pathways of adhesion formation. Second, the branched structure of the Arp2/3-nucleated actin network offers specific benefits to the assembly processes. To date, *PTK2/*FAK, *VCL/*vinculin and *FERMT2*/kindlin-2 are the sole adhesion molecules known to directly interact with the Arp2/3 complex ^29–31^. Their functions have been studied extensively on 2D substrates ^20,29–31^ but not in 3D. KOs of kindlin-2 and vinculin showed marked spreading defects in 3D collagen matrices, while FAK KO significantly increased the spreading ability of cells (Fig. 4a,b; Supplementary. Fig. 3a), which appears unsurprising given the strong upregulation of adhesions reported in FAK KO mice ^32^. Further analysis revealed that kindlin-2 KO cells also had a prominent spreading defect in 2D, while vinculin KO cells showed only a modest defect that is comparable to that of Arp3 KO cells (means 0.931 ± 0.048 *vs* 0.980 ± 0.080, respectively; Supplementary Fig. 3b,c; Fig. 1g). We therefore concluded that kindlin-2’s primary function in adhesion formation cannot be compensated in 2D and thus is likely independent of the Arp2/3-dependent adhesion formation pathways. Accordingly, we turned our attention to vinculin as its KO phenotype showed the same sensitivity to 2D vs 3D environments as that of Arp3 KOs.

**Figure 4.**
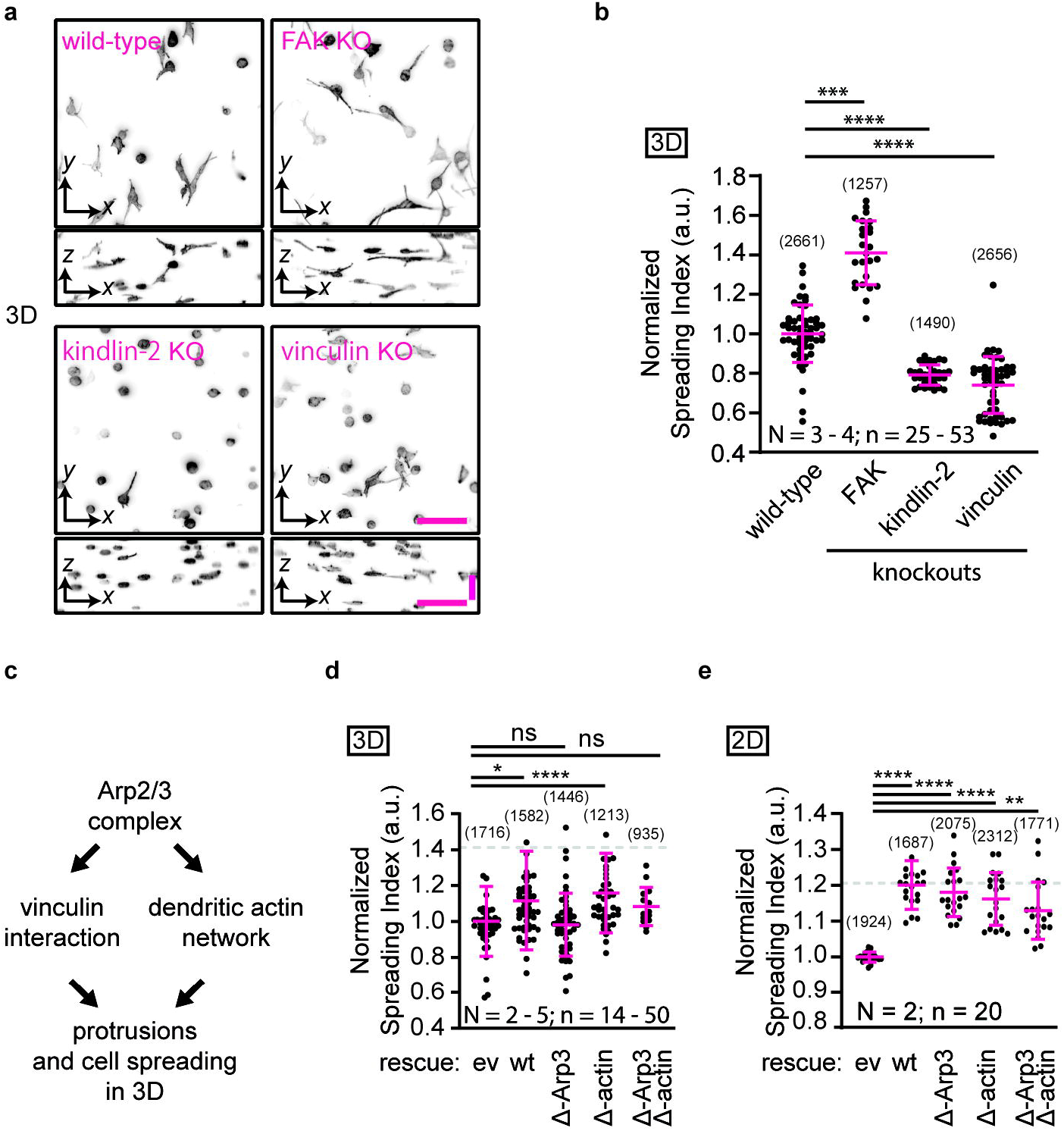
Arp2/3-vinculin interaction is required for 3D cell spreading. **a,b**, Vinculin and kindlin-2, but not FAK knockout U2OS cells have spreading defects in 3D. Cells embedded in collagen gels were stained for F-actin, imaged and quantified as described in Methods. Spreading index is normalized to wild-type control. **c**, Hypothetical model how the Arp2/3 complex contributes to cell spreading in 3D. The Arp2/3 complex can contribute to 3D cell spreading via polymerization of the dendritic actin network and/or via direct binding to vinculin. **d**, Direct Arp2/3-binding by vinculin is required for vinculin function during cell spreading in 3D. Vinculin KO cells were reconstituted with empty vector (ev), wild-type (wt), Arp3-binding mutant (Δ-Arp3), Actin-binding mutant (Δ-actin) or the combination of both mutations in full-length vinculin. Gray dotted line designates the spread levels of wild-type U2OS cells. **e**, Arp2/3-binding by vinculin is not essential for vinculin function in 2D cell spreading. Vinculin KO cells were reconstituted with empty vector (ev), wild-type (wt) or indicated vinculin mutants (Arp3-binding mutant (Δ-Arp3), Actin-binding mutant (Δ-actin)). Gray dotted line designates the spread levels of wild-type U2OS cells. All vinculin constructs were able to restore cell spreading of the vinculin KO cells comparable to that of wild-type U2OS cells. All data were collected from two to five independent experiments (N) and are shown as mean ± s.d. Each data point represents the median of the spreading indices of all cells within one image field of view (n), and normalized to ev. Total number of quantified individual cells are indicated in brackets. Kruskal-Wallis test: ns = not significant, * p < 0.05, ** p ≤ 0.01, *** p ≤ 0.001, **** p ≤ 0.0001.

Vinculin is a mechanoresponsive focal adhesion protein ^33–35^, which is required for persistent migration through 3D collagen gels in a myosin II-independent manner ^36^. The proline-rich linker region of vinculin was shown to bind and recruit the Arp2/3 complex in an actin-independent manner, and this interaction was suggested to couple adhesion formation with lamellipodial dynamics ^31^. However, this linker region has not been highlighted as a central feature of adhesion formation, as vinculin deficient of Arp2/3-binding had minimal effects on cell adhesion and spreading in 2D ^31^. Given our observation that both Arp3 and vinculin are essential for efficient spreading in 3D, we hypothesized that the Arp2/3-vinculin interaction may constitute a critical feature for cell spreading in 3D that is dispensable for spreading in 2D. To distinguish the contributions of Arp2/3-vinculin interactions from the contributions of vinculin interacting with the Arp2/3-nucleated dendritic actin networks during 3D cell spreading (Fig. 4c), we reconstituted vinculin KO cells with a) wild-type full-length vinculin, b) a vinculin mutant deficient in Arp2/3 complex-binding (P878A ^31^; Δ-Arp3), c) an actin-binding deficient mutant (I997A ^36,37^; Δ-actin), d) the combination of both mutations (Δ-Arp3 + Δ-actin), and examined their spreading in 3D relative to an empty vector control (Fig. 4d). Vinculin KO cells reconstituted with wild-type vinculin were partially rescued for their spreading defects in 3D. In contrast, expression of the Δ-Arp3 vinculin mutant failed to rescue the KO cells, whereas expression of the Δ-actin mutant rescued spreading defects to the level of cells reconstituted with wild-type vinculin. This finding is rather remarkable in view of the rich literature -- based on 2D experiments -- that defines vinculin’s actin-binding as a central feature of adhesion formation and maturation ^36–38^. The finding was corroborated by additional Δ-Arp3 mutation in the actin-binding deficient vinculin mutant, which consistently attenuated the rescue of cell spreading by the Δ-actin mutant. Most intriguingly, all mutants rescued the spreading defects of vinculin KO cells in 2D (Fig 4e). This shows that the vinculin-Arp2/3 interaction is specifically required in the 3D context, but can be compensated in 2D.

In sum, these results highlight the critical differences between cell architecture on flat, stiff 2D substrates, which has been the workhorse for high-resolution microscopy, and cell culture in 3D collagen or at least compliant 2D substrates. We show that the vinculin-Apr2/3 interaction, which is perfectly dispensable in 2D, is an essential component of cell spreading in 3D. In contrast, the vinculin-actin interaction, which is one of the primary properties of vinculin when studied in cells cultured on stiff 2D substrates, appears to play a secondary role in cell spreading in 3D. Clearly, cytoskeleton architecture and adhesions are cellular features that are particularly vulnerable to cell culture artifacts. This vulnerability is amplified by functional redundancies between ABPs, which again promotes artefactual compensation in 2D. In this case study, while the KO of Arp3 lacked a response in 2D, all studied KOs related to the Arp2/3 activating pathway – Wave2, N-WASP, cortactin – generated uniformly strong spreading deficiencies in 3D. This indicates that in a 3D scenario, the functions of Arp2/3, including the direct binding to vinculin, can no longer be compensated by other cytoskeletal components. It is thus tempting to speculate that many of the documented functional overlaps among cytoskeletal and adhesion components are merely the product of experimental limitations. As the technology allowing us to revisit some of these functions in more physiologically relevant scenarios evolves, we may find highly divergent, specific roles for many of these overlapping components.

## Acknowledgements

We thank Dr. Dick McIntosh (University of Colorado, Boulder, CO) for kindly providing U2OS osteosarcoma cells, Dr. Dana Reed (UT Southwestern Medical Center) for her logistical support and laboratory management, and Dr. Jungsik Noh for helpful discussion on functional data analysis and statistical testing. We received generous funding from the Cancer Prevention Research Institute of Texas (R1225 to G.D., RR160057 to R.F.), the National Institutes of Health (F32GM117793 to K.M.D., F32GM116370 and K99GM123221 to M.K.D., K25CA204526 to E.S.W., R33CA235254 to R.F., R01GM067230 to G.D.), and the Human Frontier Science Program (LT000954/2015 to P.R.). The content of the manuscript is solely the responsibility of the authors and does not represent the official views of the aforementioned funding agencies. This research was supported in part by the computational resources provided by the BioHPC facility located in the Lyda Hill Department of Bioinformatics, UT Southwestern Medical Center, TX.

## Author Contributions

T.I. and K.M.D. contributed equally to this work. T.I., and K.M.D conceived and designed the experiments. T.I. and J.D.C. generated cell lines, confirmed knockouts, and prepared samples. K.M.D., B.-J.C. and R.F. developed and maintained light-sheet microscopes. T.I., K.M.D. and E.S.W performed the fluorescence imaging. T.I., K.M.D., Q.S., P.R., S.J.H. and M.K.D. performed image analysis. T.I. and G.D. interpreted the results. S.P.R, N.M. and S.J.H. provided critical reagents. T.I. and G.D. wrote the manuscript with input from all authors.

## Material and Methods

### Cell Culture

Human Osteosarcoma U2OS cells were cultured in DMEM supplemented with 10% fetal bovine serum (Sigma; F0926-500ML) and maintained in a humidified incubator at 37 °C and 5% CO_2_. All cells were tested for mycoplasma using a Genlantis Mycoscope Detection Kit (MY01100). Cells were counted using Cellometer Auto 1000 Bright Field Cell Counter (Nexcelom).

### Plasmids

Annealed gene-specific single guide RNAs (gRNAs) were cloned into the BbsI site of pSpCas9(BB)-2A-GFP (PX458) ^1^. pSpCas9(BB)-2A-GFP (PX458) was a gift from Feng Zhang (Addgene plasmid # 48138). Lentiviral mNeonGreen-Paxillin-22 and mNeonGreen-Vinculin-N-21 constructs were from Allele Biotech. Vinculin mutants were generated using PCR and sequence verified with Sanger sequencing. All primers used in this study are listed in Supplementary Table 1.

### CRISPR/Cas9-mediated generation of knockout cell lines

1 × 10^5^ U2OS cells were seeded in a 6 well plate and transiently transfected overnight with plasmid PX458 containing a gene-specific sgRNA (Supplementary Table 1), together with a modified self-cleaving donor vector containing a blasticidin-resistance gene (*bsr*) cassette using Polyethylenimine (PEI). Cells transfected with SpCas9 only (no sgRNA) in combination with the *bsr* donor vector served as a negative control. The following day, the cell media was replaced. For selection, equal number of cells were seeded in a 15 cm dish in media containing 5 µg/ml blasticidine S HCl (Thermo; R21001) until the negative control was completely absent of cells. Single clonal colonies were isolated using Pyrex Cloning Cylinder (Sigma, CLS31666), and gene knockouts verified with Western Blotting. Antibodies used to verify the knockouts are listed in Supplementary Table 2.

### Two-Dimensional Cell Spreading Assay and Analysis

Collagen-coated (2 µg/ml) coverslips were placed in a 6 well plate, seeded with 1.0 × 10^5^ cells, and allowed to spread overnight. Cells were fixed with pre-warmed 4% paraformaldehyde at 37 degrees for 10 minutes. Fixed cells were permeabilized with 0.1% Triton X-100 for 10 minutes and stained with AF488-conjugated Phalloidin (Thermo; A12379, 1:40 dilution) for 1 hour. Images were acquired with an inverted Nikon Ti-E microscope at 10X magnification (Nikon 10X NA 0.25) and collected on a scientific complementary metal oxide sensor with 6.5-micron pixels, and a field of view of 2560×2160 pixels (Zyla, Andor).

For quantification, images were automatically evaluated with a program developed in MATLAB (Mathworks, Inc). Specifically, data were loaded, and foreground-background segmented using morphological opening and closing based on morphological reconstruction using a disk shaped structuring element. Foreground objects, i.e. cells, were marked, holes filled, and clusters of cells were removed based on size. All cells touching the edge of the field of view were excluded from analysis. The convex area and area were measured for each segmented object and were used to compute the Spreading Index (Convex Area divided by Segmented Area). The Spreading Index of a particular perturbation condition was normalized with the experimental control, in most cases the wild-type control cells unless otherwise indicated. Each image (field of view) instead of each individual cell was considered as a technical replicate and data point. Thus, each data point represents the median Spreading Index of dozens of cells, which also serves to reduce noise due to imperfect segmentation.

### Large Field-of-View Light-Sheet Fluorescent Microscopy (lFOV-LSFM)

For high-throughput morphological screening of knock-out cell lines, a dual-illumination light-sheet microscope was built that permits scanning of the beam in the Z-dimension as well as pivoting the light-sheet in the sample-plane to reduce shadowing and stripe artefacts ^2^. For illumination, 405 nm, 488 nm, 561 nm, and 640 nm lasers (OBIS LS and LX, Coherent, Inc.) are combined with dichroic mirrors (MUX Series, Semrock), spatially filtered and expanded with a telescope consisting of a 50 mm achromatic convex lens (AC254-050-A-ML, ThorLabs, Inc.), a 100-micron pinhole (P100H, ThorLabs, Inc.), and a 400 mm achromatic convex lens (AC254-400-A, ThorLabs, Inc.). An achromatic Galilean beam expander (GBE02-A, ThorLabs, Inc.) further increases the laser diameter 2x. All solid-state lasers are directly modulated with analog signals originating from a field-programmable gate array (PCIe-7252R, National Instruments) that have been conditioned with a scaling amplifier (SIM983 and SIM900, Stanford Research Systems).

For light-sheet generation, a 50 mm focal length cylindrical lens (ACY254-050-A, ThorLabs, Inc.) is used to focus the laser illumination into a sheet that is relayed to the 10X NA 0.28 illumination objective (M Plan Apo 10x, 378-803-3, Mitutoyo) with two mirror galvanometers (GVS001, ThorLabs, Inc.), a 50 mm achromatic convex lens (AC254-050-A, ThorLabs, Inc.), a 100 mm achromatic convex lens (AC508-100-A, ThorLabs, Inc.), and a 200 mm focal length tube lens (ITL-200, ThorLabs, Inc.). The Z-galvanometer is conjugated to the back pupil of the illumination objective, whereas the pivot galvanometer is conjugated to the sample plane. To control the light-sheet thickness, a variable slit (VA100C, ThorLabs, Inc.) was placed in the back-pupil plane of the cylindrical lens, which is conjugate to the back-pupil planes of the illumination objective.

For detection, a 16X NA 0.8 objective lens (CFI75 LWD 16XW, Nikon Instruments) and a 200 mm focal length tube lens (58-520, Edmund Optics Inc.) form the image on a sCMOS camera (ORCA-Flash4.0, Hamamatsu Photonics). A laser line filter (ZET405/488/561/640, Chroma Technology Corporation) is placed after the detection objective lens and a filter wheel (Lambda 10-B, Sutter Instrument Company) equipped with multiple bandpass filters is placed between the tube lens and the camera. The detection objective lens is mounted on a piezo-driven stage (Nano-F450, Mad-City Labs Inc.) that provides 450 µm travel range. The sample stage is a combination of a three-axis motorized stage (MP285, Sutter Instrument Company) and a rotation stage (U-651.03, Physik Instrumente). The microscope is controlled by a custom software (LabView, National Instruments).

### Sample preparation for 3D imaging

3D samples were prepared by embedding cells at a final concentration of approximately 6.7 × 10^5^ cells/ml in 3 mg/ml rat-tail collagen (Corning; 354249). 3D samples were polymerized into a custom-made 3D sample holder, by placing the holder in a 6 wells plate and at 37°C for up to 10 minutes. Once the collagen polymerized, 4 ml of media was added to the well to avoid drying of the gel, and incubated overnight in the incubator to allow cell spreading. The following day, cells were fixed with pre-warmed 4% paraformaldehyde for 30 minutes at 37 °C. Fixed gels were permeabilized with 0.1% Triton X-100 for 30 minutes, and subsequently stained with AF488-conjugated Phalloidin (Thermo; A12379, A12380).

### Three-Dimensional Cells Spreading Analysis

To evaluate cell spreading in three-dimensional environments, we imaged cells with the lFOV-LSFM and measured cell spreading using a custom image analysis script developed in MATLAB (Mathworks, Inc.). Briefly, the 0.05% brightest pixels (e.g., the mean plus four standard deviations) were used as an initial mask for the cells. Small features in the segmented objects were removed, holes in the segmented features were filled, and segmented objects touching the edge of the field of view were excluded from analysis. The Convex Volume and Segmented Volume were measured for each segmented object, and the Spreading Index was calculated by taking the ratio between Convex Volume and Segmented Volume. Normalized Spreading Index was calculated by normalizing all values with the experimental control, in most cases the wild-type control cells unless otherwise indicated. Each 3D image stack, i.e. one image field of view rather than each individual cell was considered as a technical replicate and data point. Thus, each data point represents the median Spreading Indices of dozens of cells, which aids to reduce quantification errors due to imperfect segmentation in 3D.

### Pharmacological Perturbations

The Arp2/3 Complex inhibitor CK-666 was from Millipore (182515), and was used at 250 µM. For treatment of cells in 3D, inhibitor-containing media was added to the cells as soon as the collagen gel was polymerized.

### 2D TIRF Imaging of Cell-Matrix Adhesions and Focal Adhesion Analysis

Cells were imaged with a DeltaVision OMX SR (General Electric) equipped with ring-TIRF, which mitigates laser coherence effects and provides a more homogeneous illumination field. This microscope is equipped with a 60x, NA=1.49, objective, and 3 sCMOS cameras, configured at a 95 MHz readout speed to further decrease readout noise. Images 1024×1024 pixels were acquired with an effective pixel size of 80 nm. Imaging was performed at 37°C, 5% carbon dioxide, and 70% humidity. Laser-based identification of the bottom of the substrate was performed prior to image acquisition, with a maximum number of iterations set to 10. Laser powers were decreased as much as possible (usually between 0.2-2%). Cell-matrix nascent adhesions (NAs) were detected and segmented as diffraction-limited objects, and matured focal adhesions (FA) were detected as previously described^3^. Briefly, adhesions from images with paxillin staining were segmented using a combination of Otsu and Rosin thresholds. Segmented areas larger than 0.2 μm^2^ were considered focal contacts (FCs) or FAs, based on the criteria described by Gardel et al ^4^. Individual segmentations were assessed for the area and the length, which is estimated by the length of major axis in an ellipse that fit in each FA segmentation. FA density was calculated as the number of all segmentations divided by the cell area. Nascent adhesions were detected using the point source detection used in single particle tracking ^5^. Briefly, mNeonGreen-paxillin images were filtered using a Laplacian of Gaussian filter, local maxima were detected and fitted with an isotropic Gaussian function (standard deviation: 2.1 pixel). Outliers were removed using a goodness of fit test (*p* = 0.05). NA density was defined as the number of NAs divided by the cell area derived from 5 µm from the segmented cell border. All densities were normalized to the corresponding wild-type control.

### High-Magnification Cell-Matrix Adhesion Imaging in 3D

For high-resolution adhesion imaging in 3D, a high-NA version of Axially Swept Light-Sheet Microscopy was developed ^6,7^. Illumination is provided with 445 nm, 488 nm, 514 nm, 561 nm, and 640 nm lasers (OBIS LS and LX, Coherent, Inc.), which are co-aligned with dichroic mirrors (MUX Series, Semrock), spatially filtered and expanded with a telescope consisting of a 50 mm focal length achromatic convex lens (AC254-050-A-ML, ThorLabs), a 30 µm pinhole (P30H, ThorLabs, Inc.), and a 150 mm focal length achromatic convex lens (AC254-150-A, ThorLabs). Laser polarization was controlled with a half waveplate. All solid-state lasers are directly modulated with analog signals originating from a field-programmable gate array (PCIe-7252R, National Instruments) that have been conditioned with a scaling amplifier (SIM983 and SIM900, Stanford Research Systems).

For light-sheet generation, a 50 mm focal length cylindrical lens (ACY254-050-A, ThorLabs, Inc.) is used to focus the laser illumination into a sheet. A mechanical slit (VA100C, ThorLabs, Inc.) conjugate to the back pupil of the cylindrical lens was used to adjust the effective numerical aperture of the light-sheet, and a diffraction grating at the focus on the cylindrical lens was used to create a lattice of coherent Gaussian beams. The light-sheet was relayed to the first intermediate image plane with a 75 mm achromatic doublet (AC254-050-A, ThorLabs, Inc.), a matched pair of 3 mm mirror galvanometers (6215H, Cambridge Technology) conjugate to the back pupil of the cylindrical lens, and a 60 mm focal length f-theta scan lens (S4LFT0061/065, Sill Optics). One galvo laterally scans the lattice of Gaussian beams in the X-direction and create a time-averaged light-sheet with decreased susceptibility to shadowing artifacts, and the other galvo scans the light-sheet in the Z-direction.

The first intermediate image plane was relayed to a small aperture remote-focusing mirror (PF03-03-F01, ThorLabs, Inc.) with a 100 mm achromatic doublet lens (AC254-100-A, ThorLabs, Inc.) and a 40X 0.6 NA air-immersion objective (CFI S Plan Fluor ELWD, Nikon Instruments). Prior to being focused by the objective, the light passes through a polarizing beam splitter and an achromatic quarter-wave plate, which are located in the infinity space of the objective. The mirror, located at the focus of the air-immersion objective, reflects light back through the objective, which is subsequently passed through the same achromatic quarter-wave plate, reflected by the polarizing beam splitter, and focused to the second intermediate image plane with another 100 mm achromatic doublet. Displacement of the mirror with a piezo actuator (P-603.1S2 and E-709.SRG, Physik Instrumente) results in a wavefront that deterministically scans the light-sheet along its propagation axis (e.g., the Y-dimension). Lastly, the second intermediate image plane was imaged to the sample plane with a 200 mm tube lens (ITL200, ThorLabs, Inc.) and a NA 0.71 water dipping illumination objective (54-10-7, Special Optics). For detection, a 25X NA1.1 water-dipping objective lens (CFI75 Apo LWD 25XW, Nikon Instruments) and a 500 mm achromatic doublet (49-396, Edmund Optics) creates an image which is spectrally separated with a dichroic mirror on two sCMOS cameras (ORCA-Flash4.0, Hamamatsu Photonics). A laser line filter (ZET405/488/561/640, Chroma Technology Corporation) is placed after the detection objective lens. The detection objective lens is mounted on a piezo-driven stage (P-726 PIFOC High-Load Scanner, Physik Instrumente) that provides 100 µm travel range. The sample stage is a three-axis motorized stage (Sutter Instrument, MP285), and the microscope is controlled by the custom LabVIEW software (Coleman Technologies, National Instruments).

### Cell Adhesion Characterization in 3D

Cell-matrix adhesions were detected in a fully-automated format using tools adapted from existing Danuser lab software implemented in MATLAB (Mathworks, Inc.) ^5,8^. The variation of shape and density of adhesions structures in collagen-embedded cells required the design of a detector that is flexible enough to adapt to multiple adhesion sizes and orientations, as well as sensitive enough to detect and segment the dimmest of structures. To adapt to multiple object sizes, our detector operates in a multiscale Laplacian of a Gaussian filtering framework ^9^ to estimate a scale-space signature map for each voxel and scales ranging from 100 nm to 5 microns. Excluding boundaries, local maxima in the scale-space map represents candidate adhesions locations and scale. Each scale filtering response needs to be normalized in order to be comparable and determine the most likely scale at each location. We found that normalizing by the L1 norm of the convolution filter provided best results. In order to label voxels pertaining to the background or an adhesion structure, we exploited an adaptive thresholding approach process described in our previous work for a single, diffraction limited scale ^5^. Each scale filtering step is thus combined with a convolution-based statistical test to compares the local amplitude of a hypothetical object at a given scale with the local background. In order to further separate adhesion clusters, the watershed algorithm is applied on the volume of maximum scale filtering response. For each detected adhesions, the elongation is estimated by averaging a tubularity metric measured on each voxel associated to a single adhesion. The tubularity metric is inspired by the classic vesselness estimator by Frangi and colleagues ^10^, and adapted to discriminate between flat and elongated adhesions. Let (λ_1_<λ_2_<λ_3_) be the three eigenvalues of the Hessian matrix computed at each voxel of an adhesion, we compute tubularity metric T in the range [0 … 1]:

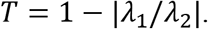

For each acquisition of a complete 3D cellular volume, the distribution of adhesion elongation per cell is estimated using a kernel smoothing function equipped with a bandwidth that is optimal under a normal distribution assumption. Due to the low count of usable volume in 3D light-sheet microscopy experiment, we visualize the uncertainty of the elongation for each condition through a random-resampling bootstrapping approach.

We assess the significance of the differences between adhesion elongation distribution in control and treated conditions by performing 100 two-sampled t-tests for 100 elongation values ranging between 0 and 1. The resulting set of p-values describes the range of elongation values where the difference between conditions is significant.

To further confirm that alleged differences between the elongation distributions of wild-type (WT) and Arp3 KO cells are significant we used the Kolmogorov-Smirnov (K-S) single-tailed test. It is well known, however, that the K-S test is very sensitive and may also detect a significant difference between two WT cells. Thus, we recalibrated the K-S test for our data by comparing the distribution of K-S statistic measured in-between individual WT cell adhesion elongation distributions to the K-S statistics measured between pairs control and Arp3 KO cells. We show that the “WT *vs* WT” group of K-S statistics is significantly lower (t-test: p < 10^−9^) than the “WT *vs* Arp3 KO” group.

