## Supplementary figures and images for "Direct Arp2/3-vinculin binding is essential for cell spreading, but only on compliant substrates and in 3D"

### Supplementary Figure 1

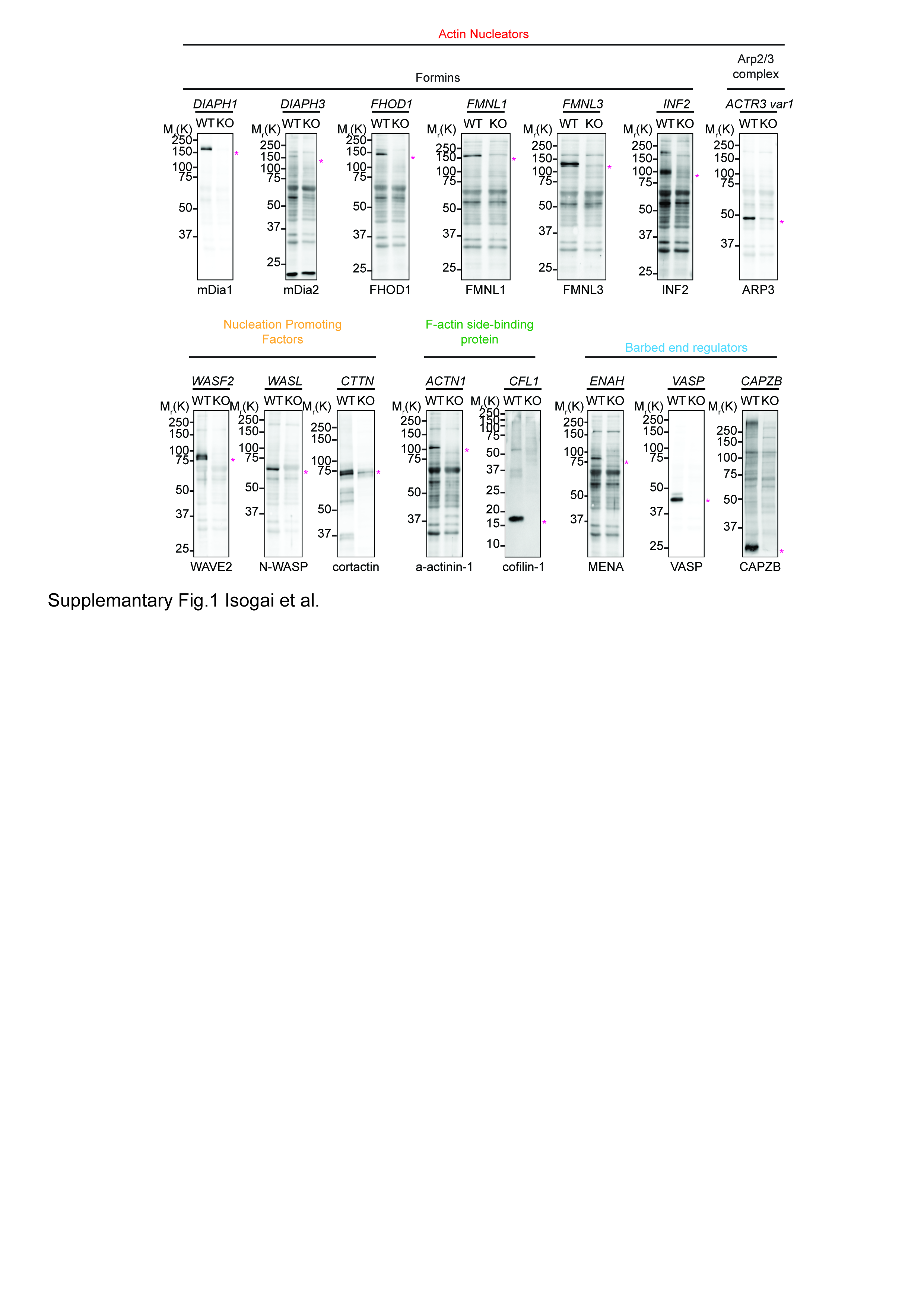

### Supplementary Figure 2

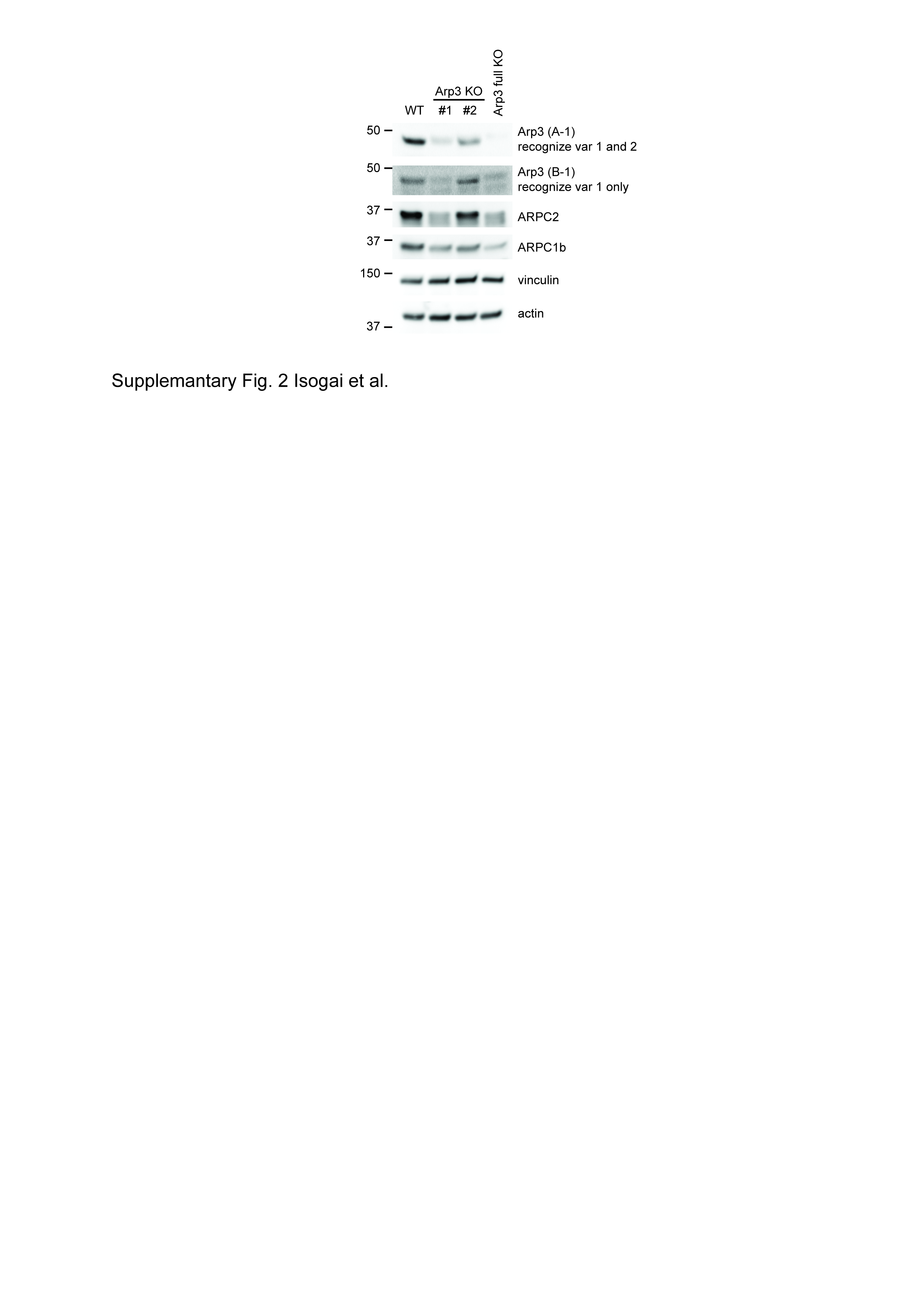

### Supplementary Figure 3

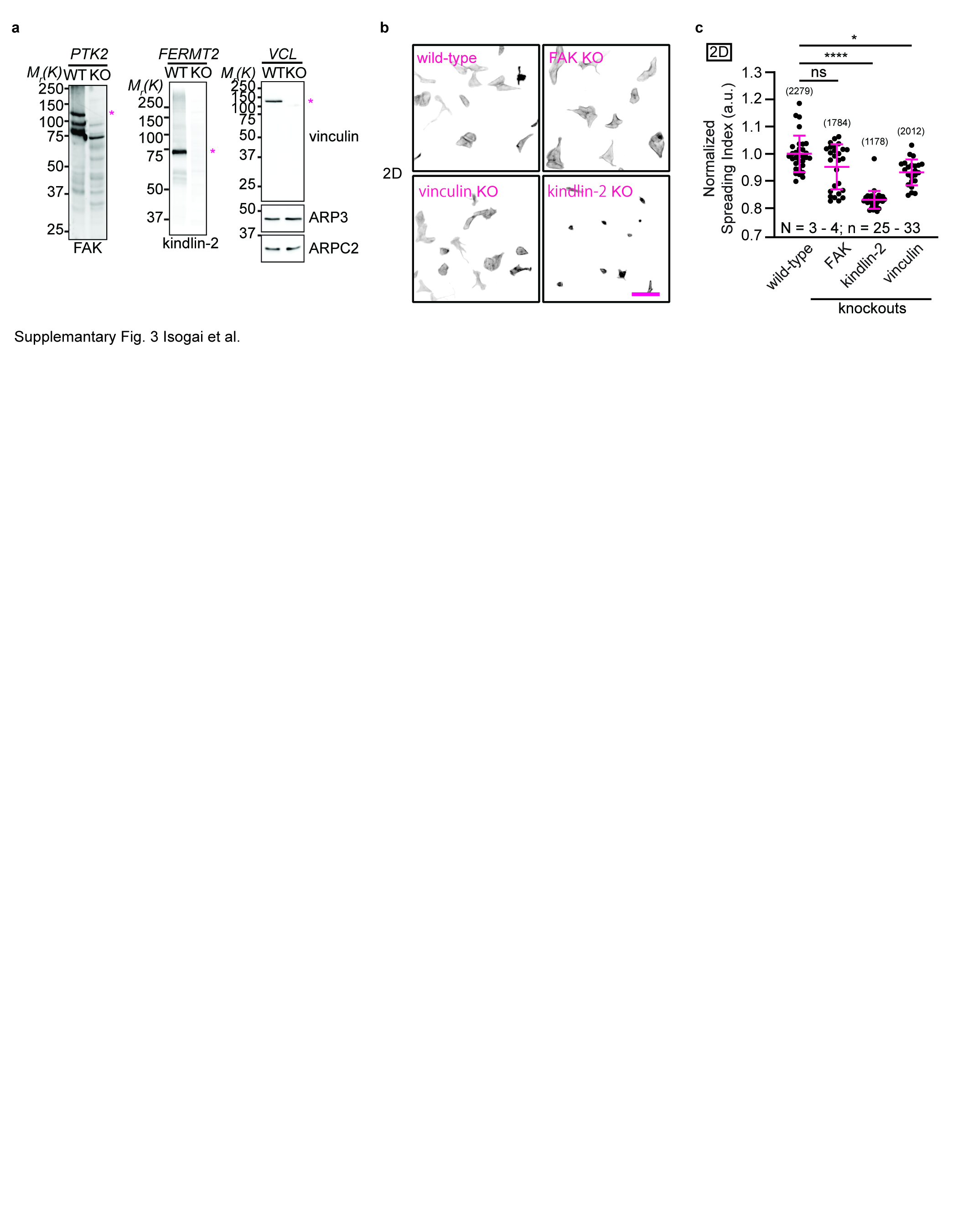
