## Supplementary Figure Legends for "Direct Arp2/3-vinculin binding is essential for cell spreading, but only on compliant substrates and in 3D"

**Supplementary Figure 1. Verification of knockout using Western Blots.** Protein lysates (30 µg) from U2OS wild-type and gene knockouts were separated by SDS-PAGE and Western Blotted for the indicated antibodies. All used antibodies and their dilutions are documented in Supplementary Table 2.

**Supplementary Figure 2. Characterization of Arp3 knockouts and evaluation of Arp2/3 complex inhibitor on 2D cell spreading.** Arp3 knockouts generated with different gRNAs were characterized with indicated antibodies. Note that Arp3 (A-1) antibody recognizes both Arp3 variants 1 and 2. Anti-Arp3 antibody (B-1) targets an epitope between amino acids 5-37 near the N-terminus of Arp3, and specifically recognizes Arp3 variant 1. Arp3 loss was associated with Arpc2, but not Arpc1b downregulation.

**Supplementary Figure 3. Effects on 2D cell spreading of KO of Arp2/3-associated adhesion molecules and rescue of vinculin KO cells. a,** Western Blot verification of FAK/*PTK2*, vinculin/*VCL*, and kindlin-2/*FERMT2* knockouts. SDS-PAGE separated protein lysates were blotted for indicated antibodies. Wild-type (WT) lysates served as control. **b,c**, Vinculin and kindlin-2, but not FAK KO results in reduced 2D cell spreading. Cells seeded on thin (2 µg/ml) collagen-coated glass were allowed to spread overnight, stained with fluorescent phalloidin, and the spreading index quantified as described in the Methods. All values were normalized to wild-type control. All data were collected from three to four independent experiments (N) and are shown as mean ±.s.d. Each data point represents the median of the spreading indices of all cells within one image field of view (n). Total number of quantified individual cells are indicated in brackets. Kruskal Wallis test: ns = not significant, * p ≤ 0.05, **** p ≤ 0.0001. Scale bar, 100 µm.
