## Supplementary Tables for "Direct Arp2/3-vinculin binding is essential for cell spreading, but only on compliant substrates and in 3D"

**Supplementary Table 1. List of Primers Used**

| Construct name | Primer 1 (5'-3') | Primer 2 (5'-3') |
| --- | --- | --- |
| pX458-*DIAPH1*-guide1 | CACCG TCTTCTTGTCCCGGGTCCCG | AAAC CGGGACCCGGGACAAGAAGA C |
| pX458-*ARP3*-KO1 guide1  (NM_005721) | CACCG ACGGCTGCCGGCCTGTGTGG | AAAC CCACACAGGCCGGCAGCCGT C |
| pX458-*ARP3*-KO2 guide2  (NM_005721) | CACCG ATTAAGGAGTCAGCAAAAGT | AAAC ACTTTTGCTGACTCCTTAAT C |
| pX458-*ARP3­­*-fullKO guide  (NM_001277140) | CACC GCTCAAAGGAGGGTGATGAA | AAAC TTCATCACCCTCCTTTGAGC |
| pX458-*CFL1*-guide1 | CACCG CGTAGGGGTCGTCGACAGTC | AAAC GACTGTCGACGACCCCTACG C |
| pX458-*ACTN1*-guide1 | CACCG TTCTGGCTGCATGTAATCGT | AAAC ACGATTACATGCAGCCAGAA C |
| pX458-*PTK2*-guide1 | CACCG TGAGTCTTAGTACTCGAATT | AAAC AATTCGAGTACTAAGACTCA C |
| pX458-*FMNL1*-guide1 | CACCG AGCTCTCCGGCCGCGGGCAT | AAAC ATGCCCGCGGCCGGAGAGCT C |
| pX458-*FMNL3*-guide1 | CACCG TGCCGGGCGGCAGCAACAAC | AAAC GTTGTTGCTGCCGCCCGGCA C |
| pX458-*DIAPH3*-guide1 | CACCG GGGAAAGCAAGATGCCGCGC | AAAC GCGCGGCATCTTGCTTTCCC C |
| pX458-*INF2*-guide1 | CACCG TCCGTGGGGTCCGAATCCTG | AAAC CAGGATTCGGACCCCACGGA C |
| pX458-*FHOD1*-guide1 | CACCG CTTGGGCGCGCAGATACCCG | AAAC CGGGTATCTGCGCGCCCAAG C |
| pX458-*VASP*-guide1 | CACCG ACGCGAAAGGAATTGGCCGT | AAAC ACGGCCAATTCCTTTCGCGT C |
| pX458-*ENAH*-guide1 | CACCG GTTCATATCTATCACCATAC | AAAC GTATGGTGATAGATATGAAC C |
| pX458-*CTTN*-guide1 | CACCG ATGACGCGGGGGCCGATGAC | AAAC GTCATCGGCCCCCGCGTCAT C |
| pX458-*CAPZB*-guide1 | CACCG ACTGTGCCTTGGACCTAATG | AAAC CATTAGGTCCAAGGCACAGT C |
| pX458-*WASF2*-guide1 | CACCG TGAGAGGGTCGACCGACTAC | AAAC GTAGTCGGTCGACCCTCTCA C |
| pX458-*WASL*-guide1 | CACCG CCGCGGAGGGTCACCAACGT | AAAC ACGTTGGTGACCCTCCGCGG C |
| pX458-*VCL*-guide1 | CACCG ATCGTGCGCGTATGAAACAC | AAAC GTGTTTCATACGCGCACGAT C |
| pX458-*FERMT2*-guide1 | CACCG CAGATGGCTGCTACGCGGAC | AAAC GTCCGCGTAGCAGCCATCTG C |
| pmNeonGreen-vinculin(P878A)-N-21 | caGAGGAAAAGGATGAAGAGTTCCCTGAGCAGAA | cTGGTGGAGGCCTAGGTGGAGGGAC |
| pmNeonGreen-vinculin(I997A)-N-21 | CCTGTCCACAGTGAAGGCCACCATGC | gcTTTGAGCTGGGTGCTTATGGTTGGGATTCG |

Gene targeting sequences for the guide RNAs are marked in red

**Supplementary Table 2. List of Antibodies Used.**

| Gene name | Antibody | Dilution |
| --- | --- | --- |
| *DIAPH1* | BD Biosciences, mouse anti-mDia1, BD 610848 | 1:1000 |
| *ARP3* | Santa Cruz, mouse anti-ARP3, (A-1): sc-48344;  Note: recognizes both variants 1 and 2 | 1:500 |
| *ARP3* | Santa Cruz, mouse anti-ARP3, (B-1): sc-374200;  Note: epitope targets variant 1 according to the manufacturer | 1:500 |
| *CFL1* | Cell Signaling Technology, rabbit anti-Cofilin-1, D3F9 XP #5175 | 1:2000 |
| *ACTN1* | Abcam, rabbit anti-alpha Actinin, ab175944 | 1:1000 |
| *PTK2* | Santa Cruz, mouse anti-FAK, (D-1): sc-271126 | 1:1000 |
| *FMNL1* | Santa Cruz, mouse anti-FMNL1, (C-5): sc-390023 | 1:1000 |
| *FMNL3* | Abcam, mouse anti-FMNL2, ab57963;  Note: also recognizes FMNL3 ^1^ | 1:1000 |
| *DIAPH3* | ProteinTech, rabbit anti-DIAPH3, 14342-1-AP | 1:1000 |
| *INF2* | Bethyl Laboratories, rabbit anti-INF2, A303-428A | 1:1000 |
| *FHOD1* | Santa Cruz, mouse anti-FHOD1, (D-6): sc-365437 | 1:1000 |
| *VASP* | BD Biosciences, mouse anti-VASP, BD 610447 | 1:1000 |
| *ENAH* | Santa Cruz, mouse anti-Mena, (21): sc-135988 | 1:500 |
| *CTTN* | Abcam, rabbit anti-Cortactin, [EP1922Y]: ab81208 | 1:1000 |
| *CAPZB* | Santa Cruz, mouse anti-CapZ-β, (52): sc-136502 | 1:1000 |
| *WASF2* | Cell Signaling Technology, rabbit anti-WAVE-2, (D2C8) XP® #3659 | 1:1000 |
| *WASL* | Cell Signaling Technology, rabbit anti-N-WASP (30D10) #4848 | 1:1000 |
| *VCL* | Santa Cruz, mouse anti-vinculin, (7F9): sc-73614 | 1:2000 |
| *FERMT2* | Millipore, mouse anti-Kindlin-2, clone 3A3: MAB2617 | 1:1000 |
